# Simple models highlight differences in the walking biomechanics of young children and adults

**DOI:** 10.1101/2021.06.17.448884

**Authors:** Vivian L. Rose, Christopher J. Arellano

## Abstract

Adults conserve metabolic energy during walking by minimizing the step-to-step transition work performed by the legs during double support and by utilizing spring-like mechanisms in their legs, but little is known as to whether children utilize these same mechanisms. To gain a better understanding, we studied how children (5-6 years) and adults modulate the mechanical and metabolic demands of walking at their preferred speed, across slow (75%), preferred (100%), and fast (125%) step frequencies. We quantified the 1) positive mass-specific work done by the trailing leg during step-to-step transitions and 2) the leg’s spring-like behavior during single support. On average, children walked with a 36% greater net cost of transport (COT; J/kg/m) than adults (*p*=0.03), yet both groups increased their net COT at varying step frequencies. After scaling for speed, children generated ∼2-fold less trailing limb positive scaled mechanical work during the step-to-step transition (*p*=0.02). Unlike adults, children did not modulate their trailing limb positive work to meet the demands of walking at 75% and 125% of their preferred step frequency. In single support, young children operated their stance limb with much greater compliance than adults 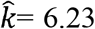 vs. 11.35; *p*=.023). Our observations suggest that the mechanics of walking in children 5-6 years are fundamentally distinct from the mechanics of walking in adults and may help to explain a child’s higher net COT. These insights have implications for the design of assistive devices for children and suggest that children cannot be simply treated as scaled down versions of adults.

## Introduction

Historically, simple models have been instrumental for understanding how humans control the lifting and forward motion of the body’s center of mass during walking. Simple models reduce the body to a point mass supported by two stiff or, alternatively, spring-like struts that characterize an “optimal” transfer or redirection of mechanical energy of the center of mass (Antoniak et al., 2019; Cavagna, GA and Margaria, R, 1966; Donelan et al., 2002; Geyer et al., 2006). With more efficient transfer or redirection, less mechanical work is required of the muscles and tendons to propel the center of mass. Lowering the mechanical work required for walking is associated with lowering the demand for consuming metabolic energy. This is now recognized as a fundamental principle of locomotion biomechanics, and one might reason that these simple models, for which these principles were derived from experimental data on adults, could be applied to young children.

However, recent insights suggest that minimizing the work performed by the legs is a dynamic process that must be learned, and which may depend on the biological and morphological constraints of the body. Bril et al. (2015) use a simple model to show how young, typically developing children (1 to 5 years) gradually learn to modulate the distance between their center of mass and center of pressure, which dynamically changes during the gait cycle. Modulating the distance between the center of mass and center of pressure is governed by the forces that the legs generate in the vertical and anterior-posterior directions. But in particular, it appears that generating forces in the anterior-poster direction (i.e. propulsive forces) is a skill learned much later in childhood, which requires tuning and control to achieve balance and elicit the desired motion in the forward direction (Brenière and Bril, 1998; Bril et al., 2015). Several studies have also highlighted differences between the typical ground reaction profiles generated by young children and adults (Dewolf et al., 2020; Takegami, et al., 1992), and have used simple models to suggest mechanical work minimization may not be the optimal strategy for young children (Usherwood et al., 2018b). This raises the question as to how the walking pattern of young children should be modeled, recognizing that that their walking mechanics may not simply be scaled down versions of adults.

For the same simple models to apply and scale to children, several assumptions must be made about their walking behavior. First, we must assume that the inertial and gravitational forces characteristic of walking will scale in proportion so that comparisons can be made across “scaled speed.” Dimensionless speed, e.g. the Froude number, is a widely used method to scale when comparing walking dynamics between children and adults; however, the Froude equation is based on the idea of dynamic similarity (R.M. Alexander and A.S. Jayes, 1983). Dynamic similarity means that multiplying all linear dimensions, time intervals, and forces by constant factors would result in identical walking patterns. Deviations from dynamic similarity are observed in young children (Kramer and Sarton-Miller, 2008; Usherwood et al., 2018b), and the Froude number makes no allowance for differences in shape. Differences in shape, or anthropomorphic proportions, may be an important consideration when making metabolic comparisons within a species, or in animals relatively close in size (Kramer and Sylvester, 2013). Alternative scaled speeds, like the Strouhal and Groucho (Alexander, 1989; Blickhan, 1989; McMahon et al., 1985) may be used when elastic forces are considered, such as in running or when a spring-like element is added to a walking template (Geyer et al., 2006).

Second, we must assume that young children have the muscular capacity to generate and control the force and mechanical work required to move the center of mass in the most economical way possible. During walking, the muscular capacity of muscles surrounding the hip and the ankle joints are particularly important for propulsion and stability (Brenière and Bril, 1998; Ishikawa et al., 2005). Compared to adults, young children display many differences in muscle and tendon properties. They have a proportionally smaller cross-sectional area of plantar flexor muscles, differences in Achilles tendon compliance (Waugh et al., 2012), slower rates of muscle force development, and lower magnitudes of relative maximum force production during isolated contractions (Radnor et al., 2018). Further experiments would help to determine how these properties influence functional or metabolic differences during walking. However, given the importance of muscles and tendons to supporting body weight and redirecting the center of mass – both critical for economic walking in adults (Donelan et al., 2002; Grabowski et al., 2005) – it is conceivable that immature muscle-tendon capacities influence a young child’s “optimal” walking solution. Of the few studies that specifically examine center of mass motion in children, normalized center of mass amplitudes in the sagittal plane are proportionally greater than in adults until around the age of 7 to 9 years (Dierick et al., 2004), which may reflect a crucial period of dynamic changes in the maturation of both muscle and tendon (Malloggi et al., 2019; Waugh et al., 2013).

Finally, we must assume that children minimize their metabolic cost during walking, and that this cost is proportional to their size. Overall, when examining walking at a range of speeds, children over 6 years exhibit an “optimal” speed that minimizes cost (DeJaeger et al., 2001). Yet, surprisingly, at “optimal” speed, the net mass-specific cost of transport (COT, J/kg/m) is up to 33% higher in children less than 9 years old (Bolster et al., 2017; DeJaeger et al., 2001; Morgan et al., 2002). With walking speeds normalized to the Froude number, DeJaeger at al. found that differences in net COT may largely be reduced, thus body size alone may account for the higher net COT observed in young children (DeJaeger et al., 2001). However, Schepens et al. found that efficiency, defined as the ratio of the total mechanical power to the net energy consumption rate, is much lower in younger children (Schepens, 2004). If mechanical power incurs a relatively greater cost in younger children, this again raises the question as to the extent to which muscular capacity and control plays a role in explaining higher net COT. Taken together, these insights suggest a functional, mechanistic explanation for why young children are less economical at walking than adults (Schepens, 2004).

To gain greater insight into a potential explanation, our objective was to use the methodology of simple biomechanical templates to investigate how the generation of limb forces and the resulting motion of the center of mass in children may differ from adults, and thus may relate to their metabolic differences. We took a systematic approach by breaking down the walking gait cycle into distinct phases of double and single support. In mature walking patterns, the legs must redirect the center of mass from a downward and forward velocity to an upward and forward velocity during double support, when both the leading and trailing limb are in contact with the ground (Kuo et al., 2005). This redirection, also called the step-to-step transition, is considered a major determinant of the metabolic cost of walking in adults (Donelan et al., 2002), and has also been studied in 12-18 month old toddlers (Hallemans et al., 2004). For our first aim, we sought to compare the positive external mechanical work (W^+^_ext_) performed by the trailing leg in young children (age 5-6 years) and adults (age 18-30), with adults representing the “ideal” behaviour. We hypothesized that 1) after accounting for differences in mass and dimensionless walking speed, the trailing limb in young children would produce less W^+^_ext_ during double support. This hypothesis was based on literature that supports the idea that during development, children gradually tune and increase their anterior-posterior propulsive forces (Bril et al., 2015), indicating that when normalized to body mass, children aged 3-8 years generate less power at the ankle than adults (Chester et al., 2006). Young children also typically produce asymmetric vertical ground reaction forces while walking (Preis et al., 1997; Takegami, M.D., 1992; Usherwood et al., 2018a), suggesting that children do not transition from one step to the next like adults.

Inspired by the bipedal walking spring mass model of Geyer et al (2006), we also set out to compare how young children and adults modulate the distance between their center of mass and center of pressure during single support, which reflects the spring-like behavior of the leg. From the perspective of a spring-mass model (Geyer et al. 2006), the redirection of the center of mass can be achieved by the release of elastic energy that was stored during single support. At the beginning of single support, when the leg extends and center of mass rises, elastic energy that was absorbed during the double support may be released, and subsequently, as the center of mass descends during the second half of single support, elastic energy can be stored in preparation for double support (Donelan et al., 2002). This appears to be the ideal behavior in adults, whereby step-to-step transitions are facilitated by the ability of the leg to store and release elastic energy and thus, operate much like a spring. In contrast to adults, a child’s leg consists of immature muscles and tendons and may not operate like an ideal spring. In addition, single support is also the phase that requires postural adjustment to stabilize the body in an upright position. The center of mass is accelerated forward and sideways at the same time toward the swing leg (Bril et al., 2015), having a destabilizing effect. Modulating the distance between the center of mass and center of pressure, i.e. the spring length, during single support may be less precise in children and require more muscle cocontraction and activation (Grosset et al., 2008; Lambertz et al., 2003). These biomechanical constraints would ultimately raise the net COT during walking and therefore, we hypothesized that 2) after considering dimensionless speed and scaling for size, the stiffness of the leg spring (*k*) in children would differ from adults.

For both of our main hypotheses, we also explored whether young children would modulate their trailing limb work and spring stiffness in the same way as adults when meeting the demands of walking at a fixed speed, but at step frequencies slower and faster than preferred. Therefore, we tested these hypotheses under conditions in which children and adults walked at their preferred speed and across a range of slow to fast step frequencies set at 75%, 100%, and 125% of their preferred step frequency. Faster step frequencies involve shorter step lengths, while slower step frequencies involve longer step lengths. At slower step frequencies, the angle between the legs at the instance of double support increases and vice versa at faster step frequencies. According to the individual limbs model, positive mechanical work at push-off depends on both the center of mass velocity and on the angle between the legs (Donelan et al., 2002). Following that model, we expected more propulsive trailing limb work at the slower step frequency condition for both adults and children. In regards to the the bipedal spring mass model, we expected that a slower step frequency would yield a decrease in the touchdown angle of the leading limb; however, both touchdown angle and *k* vary within a large range (Geyer et al., 2006), so we could not formulate a predictive hypothesis as to how *k* might change with step frequency.

Overall, we reasoned that changes in the mechanical demands of modulating step frequency would result in greater metabolic demands for young children. Specifically, when challenged to walk at the slowest step frequency, which requires the largest step lengths, young children would likely consume greater rates of metabolic energy, and hence a greater net COT when compared with the other step frequency conditions. Given known differences in a child’s muscle and tendon morphology, ankle power generation, and lower walking efficiencies, we hypothesized that when compared to adults, 3) young children would incur a greater net COT to modulate their step frequency than adults. In particular, we expected that the legs would be required to generate the greatest amount of mechanical work at the slowest step frequency condition, and that the extra mechanical demand would be more costly for children than for adults.

## Materials and Methods

### Experimental Procedures

Healthy young adults (aged 18-32 years, *n*=8) and typically developing, healthy children (aged 5-6 years, *n*=8) were recruited locally. Adult subjects gave their written informed consent to participate. Parents of child subjects gave permission and written informed consent, and children gave verbal assent to participate, in accordance with ethical guidelines and approved by University of Houston Institutional Review Board. To ensure that we captured the most natural walking patterns and comfort with the testing environment, children visited the lab for a preliminary 1.5-hour practice session the previous day. For the child cohort, a parent was present during all parts of the experiment and gave encouragement when necessary.

During the day of the experiment, subjects arrived having fasted and refrained from caffeine or exercise for at least 3 hours. Upon arrival to the lab, subjects rested for 10 min before we measured their standing metabolic rate for 5 minutes using an open circuit TrueOne 2400 metabolic system (ParvoMedics, Inc. Sandy, UT USA). The metabolic system was calibrated immediately before each session using standard gases and a 3L syringe. Following the standing trial, reflective markers (15.9 mm) were placed according to manufacturer guidelines (Lower Body Plug-in Gait, 100 Hz; Vicon 12-camera system, Nexus 1.8.5, Vicon, Oxford, UK) and the standard scaling and calibration protocol of the Nexus 1.8.5 software was followed. An additional marker was placed at the subject-specific location of the center of mass as determined by the reaction board method (Enoka, 2015). All subjects (Table 1) wore their own shirt, bike shorts, and tennis shoes with a heel-sole difference no greater than 6.4 mm.

**Table 1:** Subject Characteristics

|  | **Children** | **Adults** |
| --- | --- | --- |
| Sex (M/F) | 4/4 | 4/4 |
| Age (y) | $5.43 \pm 0.53$ | $26.38 \pm 4.40$ |
| Height (m) | $1.22 \pm 0.04$ | $1.72 \pm 0.08$ |
| Mass (kg) | $24.77 \pm 5.60$ | $75 \pm 17.40$ |
| Spring length (m) | $0.70 \pm 0.06$ | $1.03 \pm 0.06$ |
| Leg length (m) | $0.62 \pm 0.03$ | $0.92 \pm 0.05$ |
| Preferred speed (m/s) | $0.52 \pm 0.13$ | $1.13 \pm 0.18$ |
| Froude number | $0.20 \pm 0.01$ | $0.40 \pm 0.01$ |
| Step frequency achieved vs. cued (% difference) | $2.47 \pm 2.73$ | $0.66 \pm 1.30$ |

Values are mean ± SD, Percent difference in step frequency is at the preferred frequency condition

Each subject then walked on a level dual belt instrumented treadmill (1000Hz; Bertec Co. Columbus, OH USA) at their preferred speed. Preferred speed was obtained through feedback by having subjects walk at increasing increments of speed (starting from 0.3 m/s in the child group and 0.5 m/s in the adult group). Once they reached a preferred walking speed, speed was increased again and lowered if necessary, to confirm their preferred walking speed (Arellano et al., 2009). Following a 5 min rest, subjects then walked for 5 minutes at preferred walking speed and were instructed to match their step frequency to the sound of a metronome at 75, 100, and 125% of preferred step frequency (order randomized). Subjects sat for a 5 min rest period between trials. All subjects achieved a steady rate of metabolic energy consumption with respiratory exchange ratios (RER) remaining within the normal physiological range below 1.0.

### Data analysis

Walking metabolic power (W/kg) was calculated from average 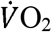 and 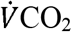 during the last 3 minutes of each trial (Brockway, 1987). The average quiet standing value was subtracted from the average walking value to yield net metabolic power (W/kg). Net metabolic power was divided by speed (m/s) to obtain net COT expressed in units of J/kg/m. Data were filtered using a 4^th^ order, zero-lag low-pass Butterworth filter with a cut-off frequency of 15Hz for force and 6Hz for kinematics. All data were processed in Matlab (R2018b, The MathWorks, Massachusetts, USA) and custom code was written to calculate values for limb work and power (Donelan et al., 2002) and for values of leg stiffness, *k*, following a spring-mass model (Geyer et al., 2006). To identify gait cycle events and periods of double and single support, we defined touchdown and toe-off as the instant when the vertical ground reaction force (GRF) crossed a threshold of 5% body weight. Double support was then defined as the portion of the gait cycle after touchdown of the leading limb, and before toe-off of the trailing limb. Single support was defined as the portion of the gait cycle between toe-off and touchdown when only one foot was in contact with the ground.

#### Individual Limbs Method

Starting from the 3 min mark of each 5 min walking trial, filtered force data from periods of double support were aggregated to determine lead leg and trail leg power and work. Following the method of Donelan et al., the velocity of the center of mass was determined by single integration. Then the external mechanical power generated by the trailing and leading limb (as shown in Fig. 2) was determined by summing together the dot product of the force and velocity of the center of mass acting in each direction (Donelan et al., 2002).

**Figure 1.**
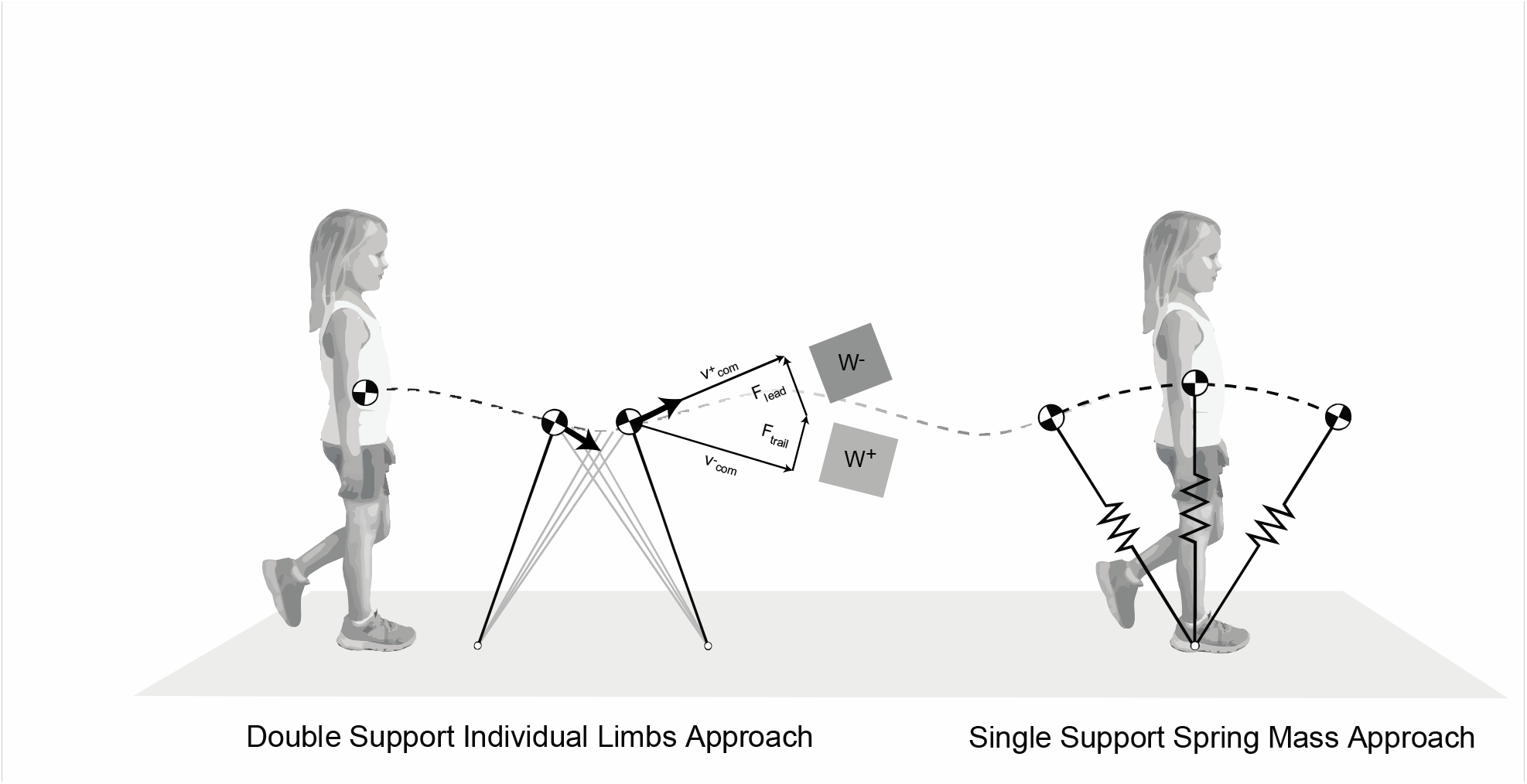
Geometric diagrams illustrate the templates used to analyze the mechanics of walking in children. For both templates, ground reaction force data were used to calculate center of mass velocity and position. Following the dynamic walking model, (left) we quantified the mechanical work generated by the individual limbs during double support (Donelan et al. 2002), a key determinant in transitioning the body’s center of mass from the trailing leg to the lead leg. In recognizing the contributions of elastic energy storage and return to the work done on the center of mass, we quantified the spring stiffness, *k*, of the leg as proposed by the spring-mass model (Geyer et al). As a simple approach, we quantified the spring-like behavior of the leg during single support, as this is the period when the leg “spring” would undergo energy release as the spring extends and the center of mass reaches it maximum height at midstance. Then, as the center of mass moves forward and its height decreases, the spring compresses and stores energy. The diagram depicting the step-to-step transition during double support is modified after Kuo et al. (2005).

**Figure 2.**
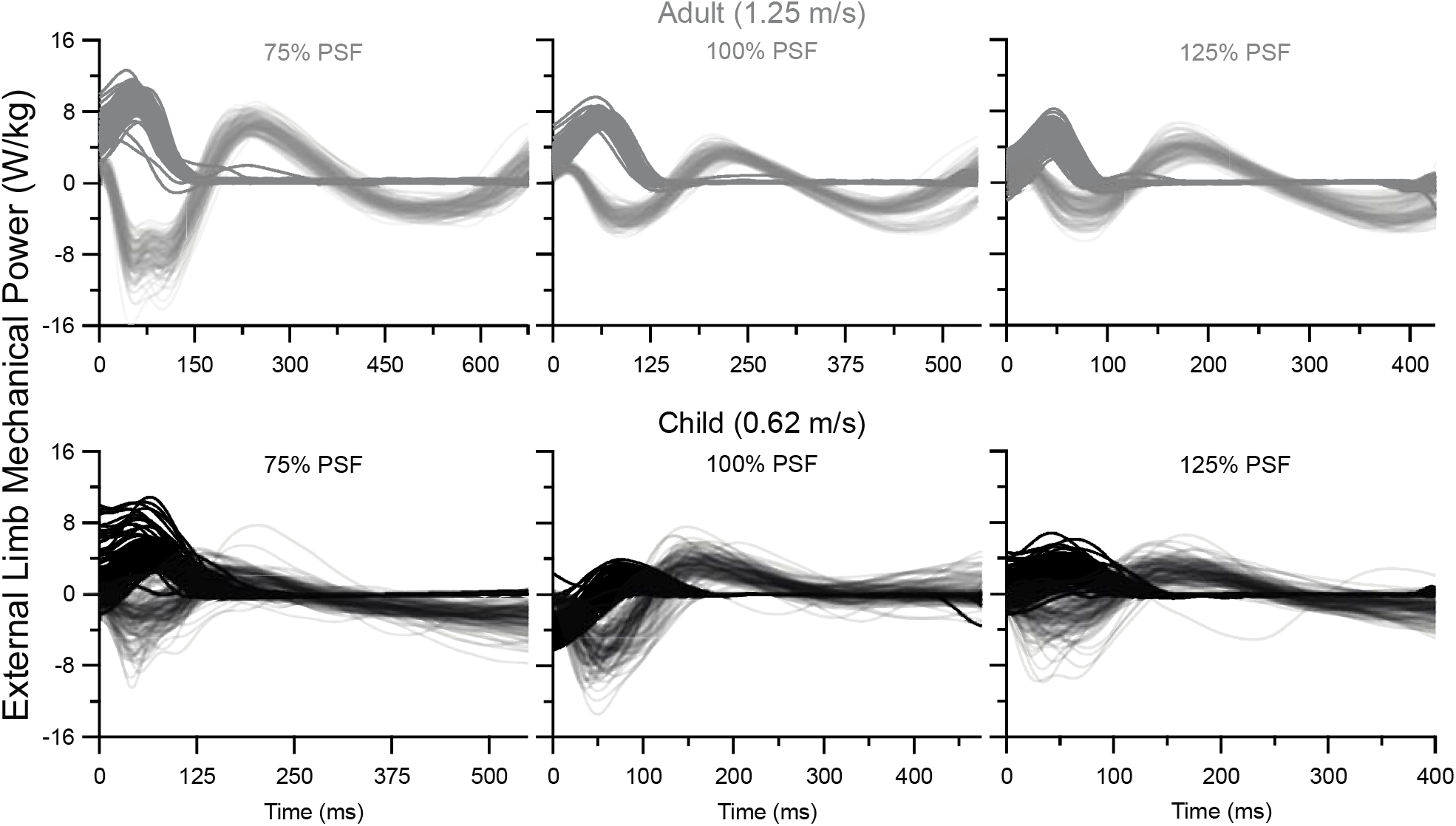
Examples of an adult (gray lines, top) and child (black lines, bottom) external mechanical power generated by the trailing (dark) and leading (light) leg during consecutive steps, under conditions of walking at their self-selected speed at 75%, 100%, and 125% of preferred step frequency (PSF). When walking at 1.25 m/s and at 100% PSF, the external mechanical power generated by the legs follows the typical pattern observed in young, healthy adults (Donelan et al. 2002), with the trailing and leading leg generating roughly equal positive and negative power during double support. In contrast, when walking at their self-selected speed of 0.62 m/s at 100% PSF, the child’s leading and trailing legs transitioned from generating negative to positive external mechanical power during double support. In general, when walking at a fixed speed, but at variable step frequencies, the adult modulated the magnitude of the positive external mechanical power generated by their trailing leg, with the slower step frequency coinciding with higher positive power and the faster step frequency coinciding with lower negative power. The child, however, exhibited a strategy whereby the trailing leg no longer generates negative and positive power during double support (as observed at 100% PSF), but primarily generates positive power when walking at the slower and faster step frequency. To highlight the greater amount of variability observed in our child group, plots of external mechanical power as a function of time consist of 100 consecutive steps.

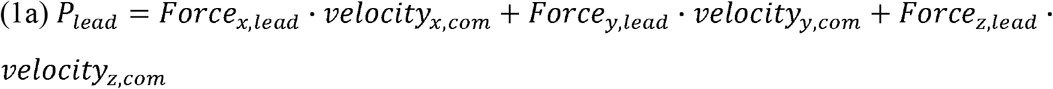

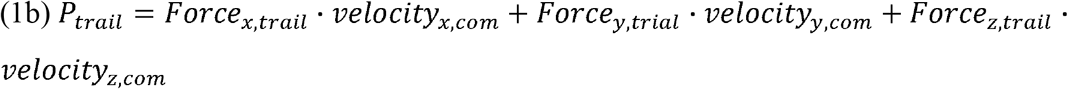

The magnitude of positive mechanical work of the trailing limb (W^+^_trail_), expressed in Joules, was then calculated as the time integral of the positive portions of trailing limb power from equation (1b). For comparisons of young children and adults at their preferred speed (m/s), the total W^+^_trail_ was normalized to mass (Schepens, 2004) and averaged across the first 100 double supports starting from the 3 min mark.

#### Single Support Spring Mass Method

Again, starting from the 3 min mark of each 5 min walking trial, filtered force data from periods of single support were aggregated. We first found the position of the center of mass using double integration and then defined a vector from the center of mass to the foot’s centre of pressure to represent a virtual 2D spring in the sagittal plane. The markers at the center of mass and pelvis were used as an offset to reflect the absolute position of the center of mass after double integration, such that the center of mass was assumed to be within the body and half the distance between the anterior superior iliac spine and posterior superior iliac spine markers. Following the work of Gard et al. (2004), we assume that this method is a close approximation to the exact location of the center of mass and more importantly, should reflect the trajectory and amplitudes that the center of mass undergoes during each step. Since we were primarily interested in the lift and propulsion of the center of mass, we reasoned that a sagittal spring was an adequate starting point for comparisons. Plotting resultant 2D GRF (N) values as a function of spring length (m) allowed us to calculate the slope of the best fit line via a least-squares regression analysis (Fig. 3). The mean of the absolute slopes estimated from the first 30 steps was reported as the value of *k*. This decision was based on our finding that adults exhibited consistent spring-like behavior that revealed best fit line R^2^ values ranging from 0.5 to 0.9. On the other hand, children exhibited spring-like behavior that was much less consistent, so for fair comparisons, we limited the R^2^ values to 0.5 which allowed us to aggregate a minimum of 30 steps.

**Figure 3.**
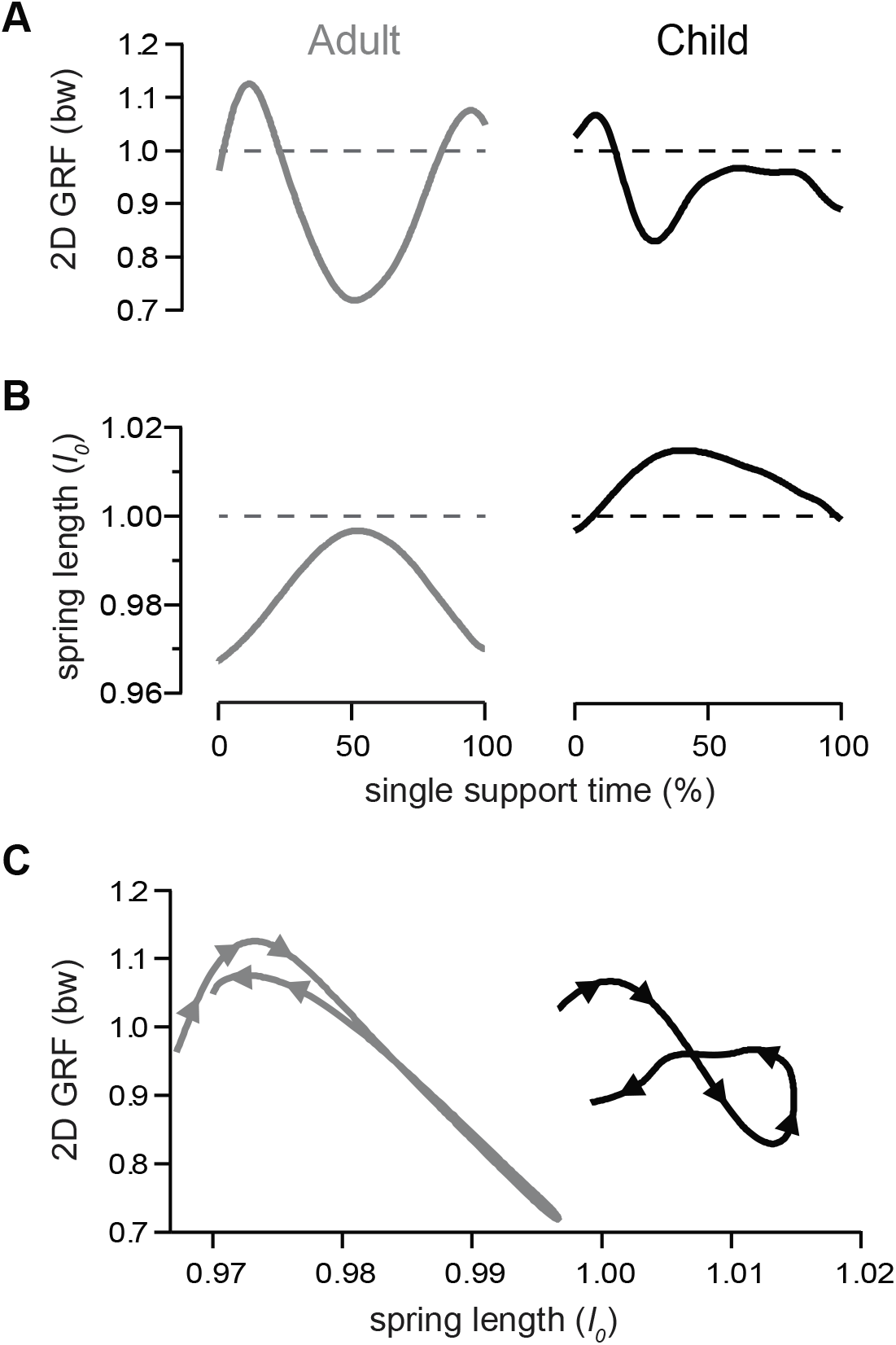
Representative adult (left, gray) and child (right, black) force and spring length curves during single support phase at preferred step frequency. (**A)** 2D resultant GRF scaled to bodyweight (bw) and **(B)** 2D spring length as measured from the COM to centre of pressure, scaled to standing height of COM (*l*_*0*_) **(C)** Force-length curves show greater asymmetry and a flatter slope for a child subject. Positive mechanical work at rebound (early single support) reflects recoil of the leg spring just after collision, while negative mechanical work in preload (late single support) reflects compression of the leg spring in preparation for propulsion. The convention here is that the absolute value of the slope equals the normalized spring stiffness,□.

#### Scaled speeds and Touchdown Angle

The Froude number, as shown in Eq. 2, normalizes speed based on pendular dynamics (R.M. Alexander and A.S. Jayes, 1983). We defined velocity, *v*, as the speed of the treadmill, and measured leg length as the distance between the anterior superior iliac spine to the distal tip of the medial malleolus. The dynamic walking model equations only apply when the body conforms to inverted pendulum dynamics (Kuo et al., 2005), so we chose to consider Froude speed in our comparison of individual limb work.

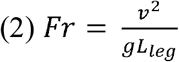

To normalize *k*, we used the Groucho speed, which was originally developed as a vertical speed parameter, combining interactions between effective vertical spring stiffness, gravity, impact velocity, and body mass (McMahon et al., 1985). It has been mathematically adapted and used in 2 dimensions by also considering leg length (Blickhan, 1989). Essentially, for both the fore-aft and vertical directions, the Groucho speed can be calculated using equation (3a) and normalized to leg length, where *v* is the resultant 2D velocity at the instant of touchdown, *g* is gravity, and ω is the natural frequency of the system as determined by spring stiffness, *k*, and body mass, *m*, as in equation (3b).

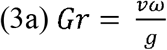

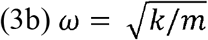

Because the Groucho number is a dimensionless speed based on a spring-mass model, we planned to use Groucho speed as a covariate in our comparison of *k*, but as explained in the Statistical analysis below, we instead used 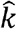, which represents its non-dimensional form. Touchdown angle was determined in the sagittal plane as the angle formed at touchdown between a line created by the center of mass and lateral malleolus marker with respect to the horizontal. We aggregated the touchdown angles from the entire 3 minutes and report the means in Table 2.

**Table 2.** Descriptive Statistics, Pairwise and Planned Comparisons

|  | Step Frequency Conditions |  |  |  |  |  |  |  |  |
| --- | --- | --- | --- | --- | --- | --- | --- | --- | --- |
|  | 75% |  |  | 100% |  |  | 125% |  |  |
|  | Mean | SD | N | Mean | SD | N | Mean | SD | N |
| $W_{\text{trail}}^+$ (J/kg)<sup>a</sup> | | | | | | | | | |
| Child | .06 | .01 | 8 | .06 | .01 | 8 | .04 | .01 | 7 |
| Adult | .16 | .01 | 8 | .11 | .01 | 8 | .09 | .01 | 8 |
| Cohen's *d* | 10.00 |  |  | 5.00 |  |  | 3.74 |  |  |
|  | ***p*<.001** |  |  | ***p*=.01** |  |  | ***p*=.03** |  |  |
| $\hat{k}$ | | | | | | | | | |
| Child | 3.57 | 3.74 | 7 | 6.23 | 5.32 | 7 | 10.66 | 4.24 | 7 |
| Adult | 6.99 | 1.29 | 7 | 10.64 | 2.33 | 7 | 16.43 | 4.09 | 7 |
| Cohen's *d* | 1.22 |  |  | 1.07 |  |  | 1.38 |  |  |
|  | ***p*=.035** |  |  | ***p*=.039** |  |  | ***p*=.024** |  |  |
| Touchdown angle (°)<sup>a</sup> |  |  |  |  |  |  |  |  |  |
| Child | 72.88 | 3.95 | 8 | 73.96 | 2.76 | 8 | 77.61 | 3.07 | 7 |
| Adult | 71.33 | 3.56 | 8 | 75.07 | 2.48 | 8 | 77.72 | 2.96 | 8 |
| Groucho/Leg length<sup>b</sup> |  |  |  |  |  |  |  |  |  |
| Child | .04 | .03 | 8 | .03 | .01 | 8 | .05 | .02 | 7 |
| Adult | .04 | .01 | 7 | .04 | .01 | 7 | .05 | .01 | 7 |
|  | *p*=.911 |  |  | *p*=.120 |  |  | *p*=.585 |  |  |

**a.** Values are evaluated at Froude 0.3 with ANCOVAs when main effect for group is significant; comparisons with □ were followed up with *a priori t*-test when significant. **b.** Independent *t*-test

**Table 3.** Metabolic R-ANCOVA results, and ANCOVA pairwise comparisons

|  | *Effects<sup>a</sup>* | *F* | *p* |
| --- | --- | --- | --- |
| Net Cost of Transport<br>(J/kg/m) | **Step frequency** | ***F*=3.805, *df* = 2,26** | **0.036\*** |
|  | **Group** | ***F*=4.003, *df* = 1,13** | **0.034\*** |
|  | Group x Step frequency | *F*=2.112, *df* = 2,26 | 0.071 |
|  | Conditions |  |  |
|  | Children and Adults at 75% | *F*=0.232, *df*=1,16 | 0.319 |
|  | **Children and Adults at 100%<sup>b</sup>** | ***F*=7.653, *df* = 1,16** | **0.008\*\*** |
|  |  | ***U* = 49** | **0.001\*\*** |
|  | **Children and Adults at 125%** | ***F*=3.752, *df* =1,16** | **0.038\*** |

**a.** Mean differences evaluated at Froude 0.3; **b.** Comparison violated assumption of homogeneity of variances, so differences analyzed with a Mann-Whitney U test; \**p* < .05, \*\**p* < .01

### Statistical analysis

We used separate mixed R-ANCOVA’s with *a priori* planned comparisons to test for differences in W^+^_trail_, non-dimensional spring stiffness 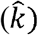, touchdown angle, and net COT. After checking for normality, we compared differences between groups at the 3 step frequencies by defining age and step frequency as a between and within subjects fixed factor, respectively. To account for differences in size and speed between children and young adults, we included in our analysis of W^+^_trail_, the dimensionless speed Froude as a covariate.

Upon initial inspection of our Groucho speeds, we removed an outlier in the adult group so statistical comparisons for *k* were based on *n*=7 in the adult group. Based on independent *t*-test at each step frequency (Table 2), we did not detect significant differences in Groucho speed between groups and therefore did not require Groucho as a covariate. We did not use Froude as a covariate for this analysis, since we were only concerned with single support and were using a theoretical model that is based on spring-mass dynamics, not pendular dynamics. To account for differences due to body size in our comparison of *k*, we transformed to non-dimensional 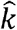 using equation 4, where *l* is the max spring length, *m* is body mass, and *g* is gravity.

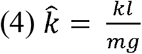

For comparisons of touchdown angle and net COT (J/kg/m), we included Froude as a covariate because these variables strongly depend on walking speed. During post-processing, we discovered that a hardware malfunction caused the force data to be erased for one child subject walking at 125% of their preferred step frequency. Therefore, statistical comparisons for mass-normalized W^+^_trail_ and 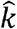 were based on a sample size of *n*=7 in the child group. For our planned comparisons, we used independent *t*-tests when normality was met or Mann-Whitney U tests when normality was not met. Finally, we plotted mass-normalized W^+^_trail,_ 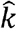, and net COT across conditions for both groups. All data was analysed in SPSS with statistical significance set at 0.05 (version 26, IBM, Armonk, NY).

## Results

### Touchdown angle

We used Froude as a covariate to compare touchdown angle between groups and confirmed that touchdown angle, an indirect gauge for detecting differences in step length, was changing with step frequency as expected. A significant main effect for step frequency (*F*_*2,24*_ =3.324, *p*=.027, η_*2*_*=*.*217*) revealed that touchdown angle increased on average from ∼72 to 78 degrees when walking from the relatively slow to fast step frequencies (Table 2). We did not detect a main effect for group or an interaction effect between group and step frequency, indicating that in response to the changing step frequencies, children and adults altered their touchdown angle in the same manner.

### Double Support Mechanics

Prior to conducting the statistical analysis, data were inspected to ensure assumptions were met for a repeated measures-ANCOVA, with Froude speed as an appropriate covariate. We detected a significant within-subjects difference of W^+^_trail_/kg across step frequencies (*F*_*2,24*_ =3.736, *p*=.039, η_*2*_*=*.*237*). As expected in adults, overall positive propulsive work was greater at longer step lengths (75% preferred step frequency) and less at shorter step lengths (125% preferred step frequency; Figure 4A). This was not so in the child group, as confirmed by our detection of an interaction effect (*F*_*2,24*_ *=5*.*216, p=*.*013*, η_*2*_*=*.*303*), indicating that children did not modulate the amount of W^+^_trail_/kg when walking at longer and shorter steps associated with the 75% and 125% step frequency condition, respectively (Figure 4A). Between group comparisons across step frequency conditions confirmed that after accounting for differences in Froude speed, children and adults generated 0.06 J/kg and 0.12 J/kg of trailing limb mass-specific positive work, respectively (*F*_*1,12*_ =14.106, *p*=.002, η_*2*_*=*.*54*). Pre-planned ANCOVA comparisons between groups confirmed that children generated less W^+^_trail_/kg than adults at each step frequency condition (Table 2, all *p* < .05).

**Figure 4.**
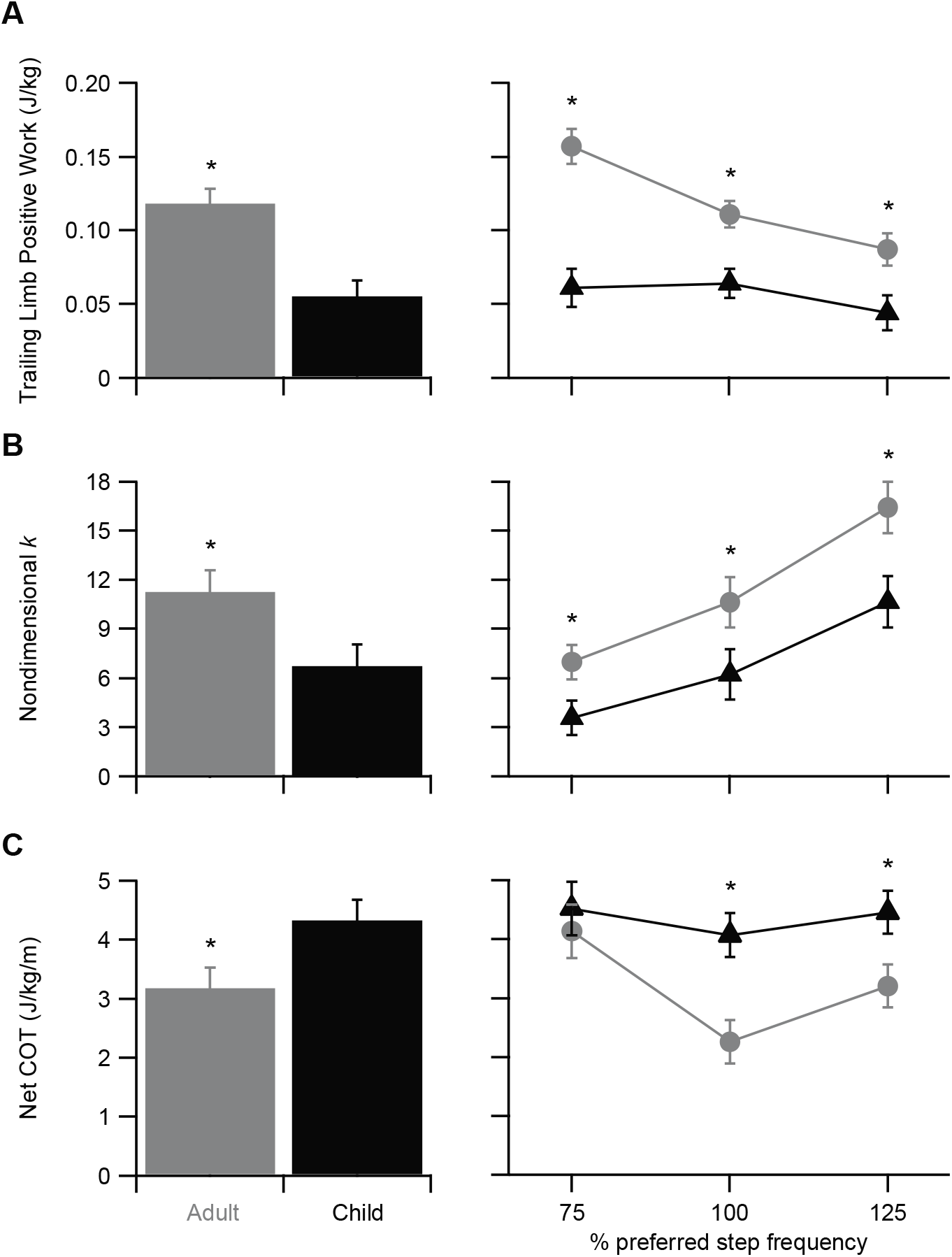
**(A)** The positive work generated by the trailing limb was smaller in children as compared to adults (left; *significant group effect *p*=0.039). In response to walking at relatively slow, preferred, and fast strep frequencies, adults (grey) altered the positive work generated by the trailing limb, however, children (black) showed little to no change in trail limb positive work (right; *significant interaction effect *p*=0.013). (**B)** When compared to adults, children operated their leg stiffness with a lower 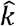. (left; *significant group effect *p* < 0.001); however, when walking across a range of relatively slow and fast step frequencies, both children and adults modulated 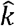. in the same way (right; no interaction effect *p* =0.41). (**C)** Differences in trailing limb positive work and 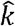. were associated with differences in the net COT required to walk. Overall, children walked with a 36% higher net COT (left; *significant group effect *p*=0.03), yet, both children and adults increased their net COT in response to walking at relatively slow and fast step frequencies, exhibiting a U-shaped trend (right; no interaction effect *p*>.05). Note that for the repeated measures ANCOVAs, average values for trail limb positive work and net COT are adjusted at a Froude number equal to 0.3. For all values, the bars represent the standard error of the mean. Significant differences between children and adults at each step frequency (right) are marked by an *, denoting *p* < 0.05.

### Single Support Spring Mass Mechanics

After checking that the statistical assumptions were met for a repeated measures ANOVA, we detected a main effect for group (*F*_*1,12*_=6.831, *p*<.001, η^*2*^ =.363) and step frequency (*F*_*2,24*_=46.43, *p*<.001, η^*2*^ =.795), but no interaction effect (*p*=0.41). At the slow step frequency associated with longer steps, 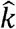 decreased, and at the fast step frequency associated with shorter steps, 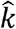 increased (Fig. 4B). Overall, children walked with an average 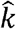 equal to 6.82, almost 2-fold more compliant than adults, who walked with an average 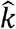 equal to 11.35. The much more compliant 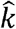 observed in children was consistent at each step frequency condition (all *p’s*<.05 for all *a priori* independent *t*-tests; Table 2).

### Net Metabolic Cost of Transpor

Prior to conducting the analysis, data were inspected to ensure assumptions were met for a mixed design and that the Froude number was an appropriate covariate. A main effect for step frequency (*F*_*2,28*_ =19.656, *p*<.001, η_*2*_*=*.*584*) revealed a higher net COT when walking at step frequencies above and below preferred. We also detected a main effect for group (*F*_*1,14*_ =31.84, *p*<.001, η_*2*_*=*.*695*), revealing that the net COT was on average, 36% higher in children (Fig. 4C). However, we did not detect an interaction effect (*p*<0.05), indicating that across slow and fast step frequencies, the net COT changed in a similar way for adults and children. And finally, when adjusted for group differences in the Froude number, the mean values show that in both children and adults, the net COT increased when walking at the relatively slow and fast step frequencies.

## Discussion

We analyzed experimental data using simple models of walking to compare center of mass mechanics and net COT in a group of young children to those of adults, who represent the ideal behavior these simple models are based upon. One key finding was that trailing limb positive work, W^+^_trail_/kg, was significantly lower across conditions in children. This supported our first hypothesis. However, in contrast to the other variables tested, the magnitude of this difference changed depending on the step frequency condition. The largest deficit was evident at the 75% step frequency—the condition that required subjects to take the longest steps. From Donelan et al., (2002) and Kuo et al. (2005), it is predicted that longer steps will increase the collision of the lead leg with the ground, effectively increasing negative work generated by the leading limb. To account for this greater negative work, the trailing limb must generate more positive work. Yet children did <u>not</u> generate the positive work that would be expected to account for the increased collision forces that are associated with taking long steps at a fixed speed.

When walking at the preferred step frequency condition, we found that on average, the trailing limb in children is used to both absorb and generate work, while in adults, the trailing limb only generates positive work, providing almost 100% of all their propulsive positive work (e.g., see Fig. 2, middle column). In adults, there seems to be a clear role for the trailing limb to generate positive work and the leading limb to generate negative work, whereas young children used both limbs to generate both positive and negative work. Halleman’s et al. (2004) reports that in toddlers (12-18 months), an inefficient inverted pendular mechanism of energy exchange contributes to differences in external mechanical work. Toddlers are described as utilizing a “tossing gait” where work that is performed to lift the center of mass against gravity is much greater than work that is performed to propel the center of mass. In our age group of 5-6 years, gait patterns are considered more mature than in toddlers. Yet, when partitioning the positive and negative work generated by each limb double support, we also found (as in Hallemans et al.) that our child subjects tended to generate more mass-specific work to lift the center of mass than in adults. However, one key difference was that the mass-specific work to lift versus to propel the center of mass was approximately equal for our child subjects (Fig. 5), suggesting a possible shift with age toward more mature patterns when work to propel the center of mass dominates.

**Figure 5.**
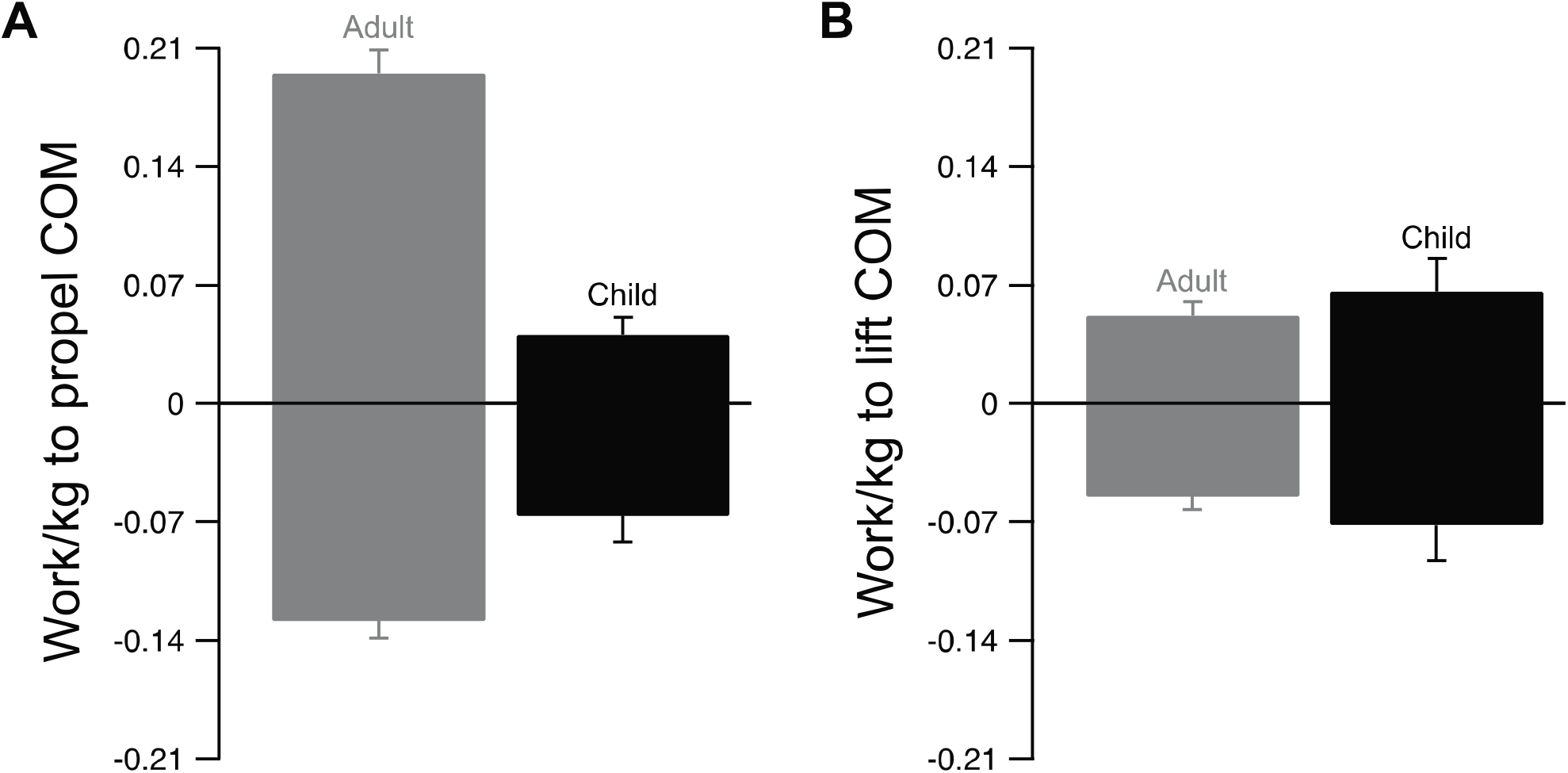
Partitioning the mean mass-specific work of both limbs reveal that children performed **(A)** much less work to propel the COM than adults, but **(B)** slightly higher work to lift the COM. As noted in the textNote that the trailing limb in the child group performed both positive and negative work in double support, while the trail limb in the adult group was only used to generate positive work.

In single support, we found that after scaling for size and speed, 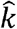 is approximately 40% lower in our child group, which supported our second hypothesis. While 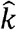 was substantially lower in our child group, they did modulate 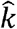 in the same way as adults. As shown in Fig. 4B, both young children and adults increased 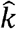 when meeting the mechanical demands of walking at fixed speed, but at relatively slower and faster step frequencies. The tendency of children to modulate 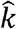 was independent of changes in the amount of positive work generated by their trailing limb during double support. This behavior deviated from that observed in adults, where the positive work generated by the trailing limb during double support decreased and 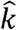 increased across slow to fast step frequencies. Unlike trailing limb work, which has been shown to rely primarily on plantar flexor muscles and tendons (Fukunaga et al., 2001; Ishikawa et al., 2005; Sawicki and Khan, 2016), 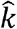 can be influenced by multiple muscles and joints, such as the ankle, knee, and hip. Thus, it is possible that both the more compliant 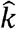 and inability to generate as much scaled work and power by the trailing limb may be related to a young child’s developing muscular capacity. Over time, as children learn to coordinate their leg and hip muscles to generate propulsive work, the motion of the center of mass during walking may become smoother, reflecting patterns that are observed in adults. Given evidence of structural differences of immature plantar flexor muscle and tendon (Radnor et al., 2018; Waugh et al., 2012) and functional differences in muscle and tendon interaction in children measured directly by ultrasound while hopping (Waugh et al., 2017), it seems likely that the coordination of the plantar flexor muscle-tendon mechanics responsible for efficient propulsion and redirection of the center of mass in adults is not fully formed and learned at 5-6 years of age.

The unequal contribution of work by the trailing limb in young children could also help explain their decreased efficiency (Schepens et al 2004), since positive work is necessary to restore the energy that is lost from the unavoidable collision phase of double support (Donelan et al., 2002). Lost energy must then be replaced with more costly compensation strategies, possibly at other joints such as the hip (Lye et al., 2016; Sawicki et al., 2009), and with coactivation of other muscles (Lambertz et al., 2003). This would ultimately raise the net COT, which is what we observed in our 5–6-year-old child group. Consistent with previous studies (DeJaeger et al., 2001; Morgan et al., 2002), our children had a higher net COT at each step frequency condition, even after accounting for differences in Froude speed. When compared to adults, however, the net COT in children did not fluctuate much in response to changes in step frequency (Figure 4C). Thus, we reject our third hypothesis that when compared to adults, walking at the lowest step frequency condition would be relatively more costly for children.

During the first 7-13 years of life, walking patterns vary in their maturity, so interpretation of our results is limited to children aged 5-6 years. At this age, we observed a high amount of variability in the ground reaction force patterns, consistent with previous studies (Kraan et al., 2017). This variability ultimately meant that we were only able to reliably take 30 steps from each trial that met our condition of a reasonable linear fit between force and spring length changes during single support. It is possible that the variability in ground reaction force patterns might have another purpose, such as for motor learning, where the goal is to learn how to efficiently redirect the center of mass. Indeed, the complex relationships between age, size, mechanics, and the spatiotemporal components of walking remain to be understood, and we lack a complete understanding of their contributions to the higher net COT observed in children. Our results should be interpreted with caution because the sample size was small, though it is worth noting that our effect sizes (Cohen’s *d*) are between 1.07-10 (Table 2), which may be helpful for future reference.

In summary, when compared to adults, we found that when walking at a fixed speed, but at relatively slow and fast step frequencies, 5–6-year-old children generated significantly less positive work by their trailing limb during double support. We also found that 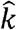, the spring-like behavior of the leg, was much more complaint in children than adults. These variables were scaled to speed, mass, and leg length, yet the mechanics of walking in children departed substantially from adults, who have long been seen as the ideal behavior for economical walking. Altogether, our findings suggest that the simple models used here cannot be scaled down and used to adequately characterize a child’s walking mechanics and energetics. The implications from this work are worth further consideration because simple models form the basis for more complex theoretical and computational models and aid in the design of assistive devices such as lower limb prostheses and orthoses (Delp et al., 2007; Gard and Childress, 2001; Geyer and Herr, 2010). As opposed to relying on models that are scaled down versions of adults, we propose that a limb-level biomechanical analysis of walking in children may facilitate the selection and tuning of assistive devices that are specifically designed for children. It is well known that adults can successfully tune and utilize the Achilles tendon to store and release energy to generate propulsive work (Fukunaga et al., 2001; Ishikawa et al., 2005; Lichtwark and Wilson, 2008), directing the attention of engineers and clinicians to the ankle as a first line target for assistive technology in gait rehabilitation. However, little is understood about plantar flexor muscle-tendon function in children, who also require this technology and, in our study, did not generate comparable positive propulsive work by their trailing limb. Combining a limb-level biomechanical analyses with ultrasound measurements of plantar flexor muscle-tendon function is of future interest, as this would help understand the unique strategies that children use to meet the mechanical power demands of walking across a range of speeds and uphill/downhill conditions.

## Acknowledgements

We thank the children and their parents for their participation in this study. We acknowledge Anna Larsson, Daisey Vega, and Danny Guevara for their assistance with data collection and for UH department of Health and Human Performance for supporting the completion of this study and manuscript preparation.

## Author Contributions

Conceptualization: V.L.R., C.J.A.; Methodology: V.L.R., C.J.A.; Software: V.L.R., C.J.A.; Validation: V.L.R., C.J.A.; Formal analysis: V.L.R., C.J.A.; Investigation: V.L.R; Resources: C.J.A.; Data curation: V.L.R., C.J.A.; Writing -original draft: V.L.R; Writing -review & editing: V.L.R., C.J.A.; Visualization: V.L.R, C.J.A.; Supervision: C.J.A.; Project administration: C.J.A.; Funding acquisition: C.J.A.

